# Mesoscale imaging of the human cerebellum reveals converging regional specialization of its morphology, vasculature and cytoarchitecture

**DOI:** 10.1101/2025.05.23.655715

**Authors:** N Priovoulos, PL Bazin, EJP Brouwer, JF Mejias, MHS de Buck, A Alkemade, W van der Zwaag, MWA Caan

**Affiliations:** Department of Biomedical Engineering and Physics, Amsterdam University Medical Centers, University of Amsterdam, Amsterdam, The Netherlands; Spinoza Centre for Neuroimaging, Royal Netherlands Academy for Arts and Sciences, Amsterdam, The Netherlands; Oxford University Centre for Integrative Neuroimaging, FMRIB, Nuffield Department of Clinical Neurosciences, University of Oxford, Oxford, UK; Full Brain Picture Analytics, Leiden, The Netherlands; Computational and Cognitive Neuroscience and Neuroimaging, Netherlands Institute for Neuroscience, Royal Netherlands Academy for Arts and Sciences, Amsterdam, The Netherlands; Cognitive and Systems Neuroscience Group, Swammerdam Institute for Life Sciences, Faculty of Science, University of Amsterdam, Amsterdam, The Netherlands; Research Priority Area Amsterdam Brain and Cognition, University of Amsterdam, Amsterdam, The Netherlands; Integrative Model-Based Cognitive Neuroscience Unit, Department of Psychology, University of Amsterdam, Amsterdam, The Netherlands

## Abstract

The human cerebellar cortex contains the highest density and number of neurons in the human brain, yet this thin and tightly-folded structure has remained largely inaccessible to in-vivo imaging. We introduce an imaging framework to enable high-resolution, comprehensive imaging of the human cerebellar cortex. We validated the in-vivo estimates of cortical morphology against post-mortem measures. Crucially, our findings challenge the commonly held view of the cerebellar cortex as being uniform. Our findings reveal interlobular heterogeneity in both cortical morphology and vascular organization. We demonstrate that this spatial heterogeneity correlates with granular layer cell density. This convergence of morphology, cytoarchitecture and vascularization offers new mechanistic insights and reframes how cerebellar structure and function should be interpreted in health and disease.

## Introduction

The cerebellar cortex has greatly expanded throughout primate phylogeny^1,2^, containing the majority of the neurons of the human brain, densely packed in a distinct laminar architecture^3^. The emergence of such a large and complex structure challenges the long-held view that the cerebellar cortex serves a singular computational role, rooted in its seemingly uniform microstructural and circuitry substrate compared to the neocortex^4,5^. Although the notion of a uniform cerebellar cortex is evolving^6^, with transcriptomic and functional regional differences becoming apparent^7^, research on local cerebellar specialization remains largely confined to animal studies: accurately reconstructing the thin and intricately-folded human cerebellar cortex is beyond the reach of most modern 2D and 3D acquisition and analysis techniques even post-mortem^8,9^, let alone in-vivo. However, given the sheer size of the human cerebellar cortex^9^ and its broad involvement across human cognitive domains^4,10–12^, understanding the cerebellar cortical substrate in humans is crucial.

In the neocortex, functional and cytoarchitectural specialization was obvious to the pioneers of neuroscience^13,14^. The development of MRI and matching analysis tools^15^ enabled non-invasively exploring this specialization: the 3D segmentation and reconstruction of the neocortical surface helped derive regional morphological measures in-vivo (e.g. cortical thickness). These measures were shown to be sensitive to the underlying neuronal density, laminar cytoarchitecture and interregional connectivity^16–18^. Neuronal subtypes, density and connectivity were likewise linked to energy demands and vascularization^19–21^.

These relationships between neuroimaging-derived measures and underlying cytoarchitecture have become central to human neuroscience, enabling insights into genetic^22–24^, development-^25–27^ and disease-related^28,29^ effects and promising new disease markers.

By comparison, the cerebellum remains underexplored: current in-vivo neuroimaging measures capture only gross anatomical structures, lacking the resolution to resolve cortical details: this is because the cerebellar cortex is thinner (0.8-1.2mm)^8^ than the neocortical (2-4mm), highly-foliated and tightly-packed (0.2-0.3mm between banks)^9^, placing it beyond the reach of standard neuroimaging spatial resolution. As a pragmatic approach, in-vivo neuroscientific and clinical research often relies on coarse approximations of the cerebellar structure^30^. Consequently, the crucial work of linking in-vivo neuroimaging measures with the underlying cytoarchitecture remains unaddressed, impeding the interpretation of human neuroscientific research in the cerebellum. Post-mortem research suffers from similar limitations: the intense cerebellar foliation necessitates 3D approaches to explore regional differences.

In this study, we explored the relationship between in-vivo cerebellar cortical morphology and vascularization and post-mortem morphological and cytoarchitectural features. To achieve this, we developed an imaging framework to capture the entire human cerebellum in-vivo in 3D and high spatial resolution, by integrating ultra-high-field MRI with optimized radiofrequency transmission and easy-to-use prospective motion correction.

Ultra-high field MRI greatly facilitates high-resolution imaging due to an increased signal-to-noise ratio compared to typical field strengths. We further built a segmentation pipeline to automatically segment the cerebellar cortex across in-vivo MRI, post-mortem MRI and 3D-reconstructed histology while retaining cerebellar cortical details. These allowed us to quantify the in-vivo regional cerebellar morphology and vascularization in 3D and compare with the cytoarchitecture.

## Results

### In-vivo cerebellar imaging framework

To image the cerebellar cortex, we scanned 10 participants with a 0.4mm isotropic Magnetization Prepared 2 Rapid Acquisition Gradient Echo scan (MP2RAGE; voxel-volume=0.064mm^3^) in a B_0_=7T human-bore MRI scanner (Philips Achieva). The MP2RAGE technique achieves white and gray-matter T_1_-weighted contrast while inherently reducing inhomogeneities relating to the receive and transmit fields^31^. To retain the nominal 0.4mm resolution in the face of involuntary participant motion, we implemented a prospective motion-correction technique, as we have published before^30,32,33^. This was achieved by interleaving quickly-sampled, low-resolution (2mm isotropic) 3D images of the fat around the skull in-between acquisition segments of the higher-resolution 0.4mm scan (Fig.1A)^34^.

**Figure 1:**
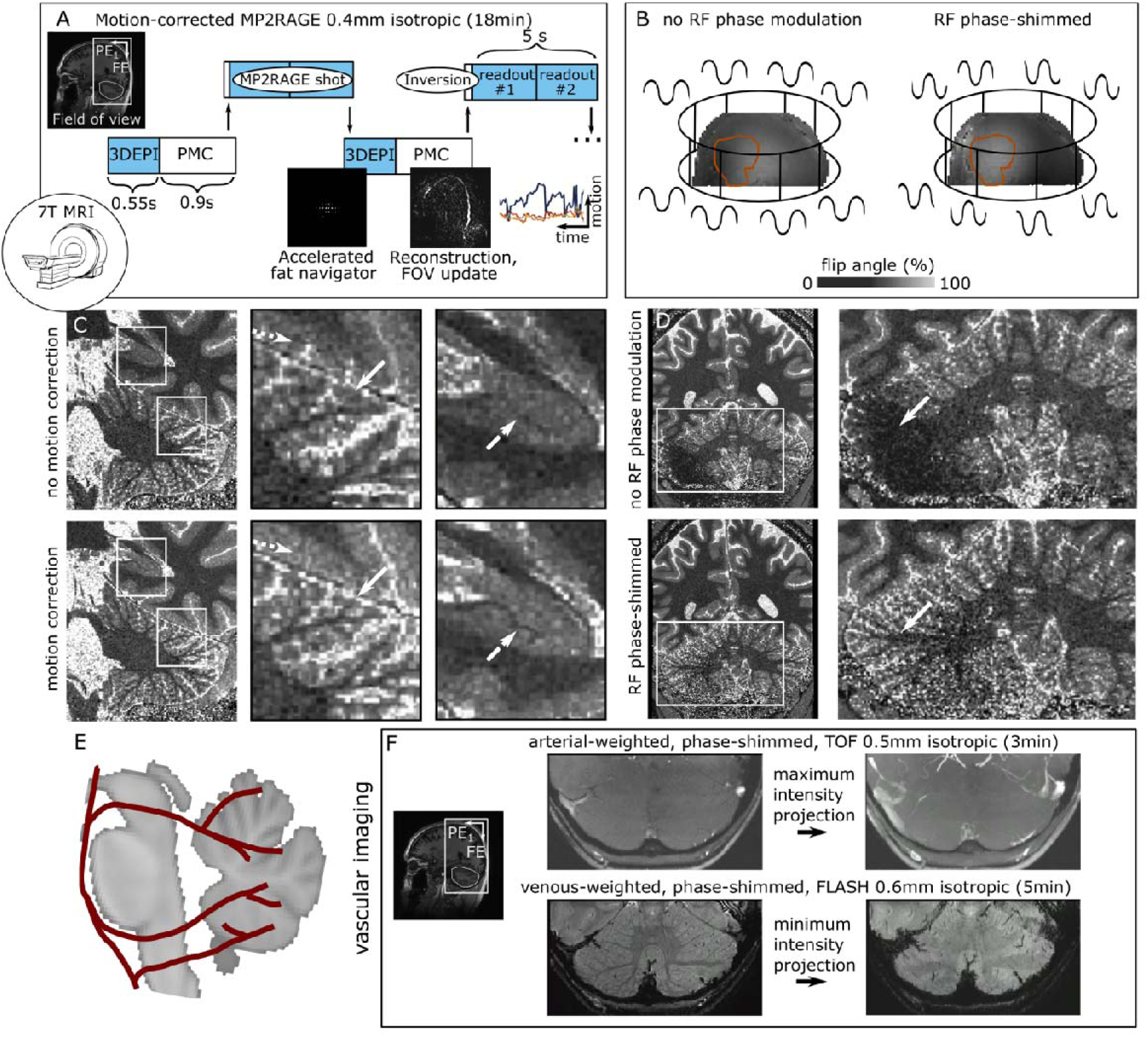
Human cerebellum imaging framework. A, in a B_0_=7T MRI human-size scanner, we implemented prospective motion correction (PMC), based on rapidly-acquired fat-navigators and update of the field of view (FOV) between successive MP2RAGE shots. B, Motion correction was combined with a phase modulation of individual radiofrequency (RF) transmission channels (right) to improve signal in the area of the cerebellum (red ROI). C, Images with and without motion-correction and close-ups in one individual. D, Images with and without RF phase shim. E, Sketch of cerebellar arterial microvasculature. F, Example time-of-flights (TOF) and venograms and matching maximum/minimum intensity projections for arterial and vascular visualization.

The human cerebellum is outside the area that MRI transmit coils are optimized for, resulting in signal drop-off with typical whole-head 7T coils^35^. To achieve homogeneous excitation in the cerebellum without increasing scan time, we applied phase offsets in each transmit channel of our parallel -transmit head coil (8Tx/32Rx Nova Medical)^36^. These phase-offsets were optimized previously over a separate participant group, specifically improving the signal in the cerebellum (Fig.1B). In combination, this motion-corrected, parallel-transmit, ultra-high field MRI imaging framework allowed us to maximize the resolution benefits from imaging at B_0_=7T while attenuating motion and transmit artifacts in the cerebellum with minimal increase in user load. The added value of our imaging approach is illustrated visually in acquisitions with and without motion correction and with and without cerebellar-optimized transmit phase-offsets (Fig.1C and Fig.1D respectively).

To comprehensively image the cerebellar vasculature, we implemented two scans: a single-slab, phase-shimmed 3D time-of-flight magnetic resonance angiography (TOF-MRA) scan weighted towards the cerebellar arteries and a 3D susceptibility-weighted, phase-shimmed venogram weighted towards the cerebellar veins. The sagittal field-of-view was set parallel to the brainstem to minimize upstream saturation of inflowing blood and thus enhance TOF-MRA vessel contrast (Fig.1E). The TOFs and venograms showed the expected image contrast, as can be demonstrated from example maximum-intensity projections to visualize the arteries (Fig.1F, top) and minimum-intensity projections to visualize the veins (Fig.1F, bottom).

### Automated segmentation of the human cerebellar cortex across modalities

Similar to the cerebral cortex, processing of the cerebellar images requires the delineation of the areas of interest, i.e. the white and gray matter. Compared to the cerebral cortex, however, extracting detailed and topology-preserving segmentations is challenging, due to the highly-foliated and tightly-packed cortex^9^. Adjacent structures, like the dura and sinuses can have similar MRI signal intensity and spatial-frequency characteristics to the cerebellar cortex, further confounding segmentation^37^. The detailed segmentation of the cerebellar cortex to the level of individual folia has been achieved only recently and post-mortem, following either 12h MRI scan time for n=1 human brain and subsequent laborious manual effort^9^ or 3D-reconstructed histology (BigBrain data^8,38^). Due to the lack of in-vivo data with sufficiently high spatial resolution^39^, automated segmentation approaches that resolve fine cortical details (similar to what is currently available for the neocortex^15^) have not been published yet: current tools are pragmatically constrained to the main branches of the cerebellar cortex^40^.

Here, we combined an atlas and intensity-based segmentation approach to automatically segment the cerebellar cortex using Nighres 1.4.0^41^. We first denoised the MP2RAGE images using local complex-valued PCA denoising^42^ followed by an initial atlas-based classification of the cerebellum. The cerebellar white and gray matter and surrounding cerebrospinal fluid were segmented by combining fuzzy c-means classification with 2D-ridge detection to extract the high spatial frequency details of the cerebellar cortex. A distance function from the cerebellar surface was employed to differentiate the dura from white matter. A distance function from the white matter surface was used to probabilistically limit the growth of gray matter into sinuses and retain the fine cortical detail. A topology-correction step was employed to preserve the spherical topology of the cerebellar cortical surface^43^ (S-Fig.1A).

For validation, we compared the automated segmentation with semi-manual segmentation (approach described here:^30^) in the same datasets. The derived cortical surface from the automated algorithm was visibly less sensitive to noise, while still differentiating small cortical branches (Fig.2A-C, S-Fig.1B). As additional validation, we implemented the segmentation for typical spatial resolutions achieved at B_0_=3T and B_0_=7T MRI. The resulting segmentations consistently differentiated cortical branches across this spatial resolution range (S-Fig.1C), suggesting that our segmentation approach can be applied in conventional imaging conditions.

**Figure 2:**
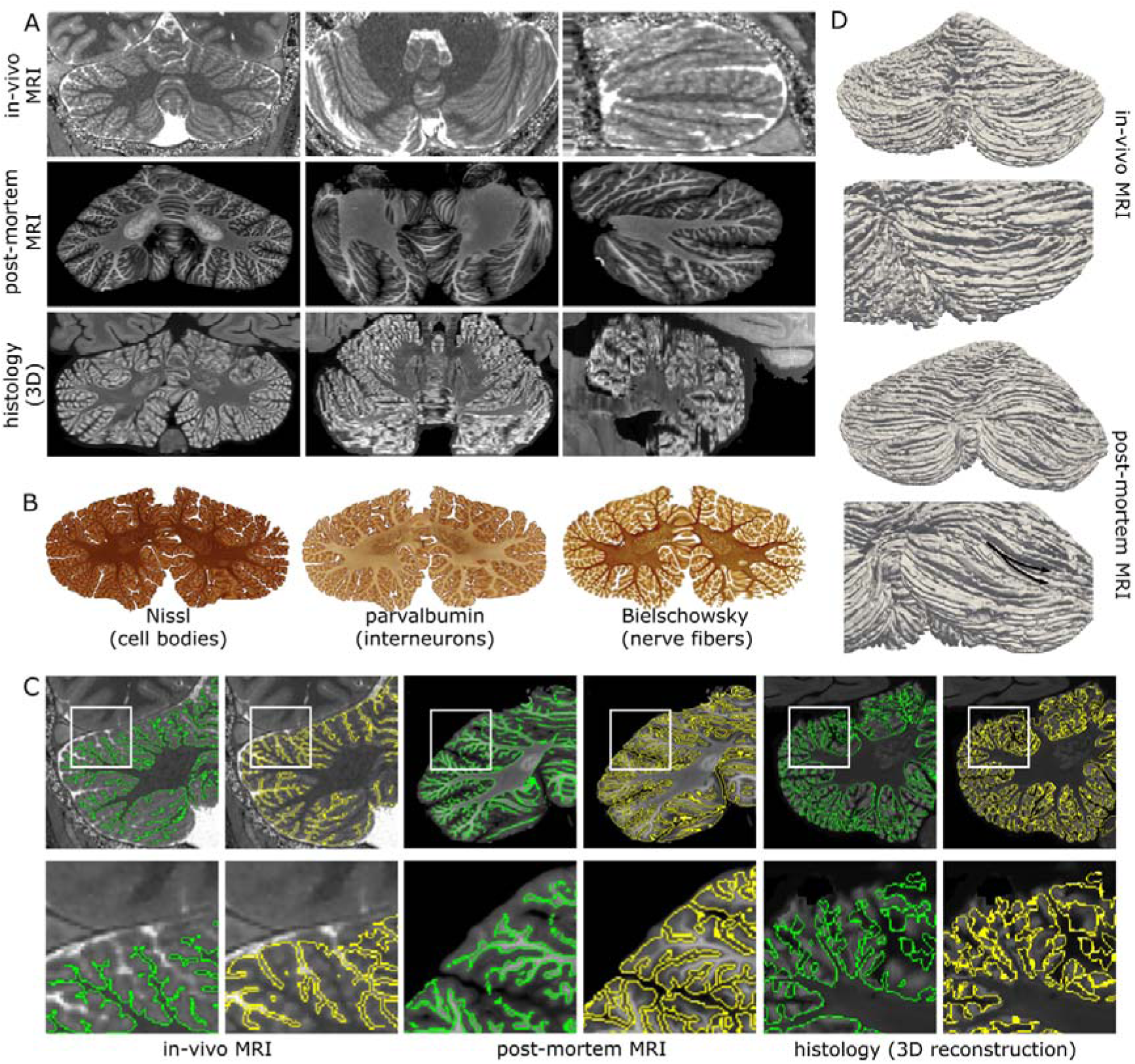
A, Example data from in-vivo MP2RAGE (left), postmortem MRI (middle) and 3D reconstructed histology. B, Examined immunohistology stains C, Example cerebellar segmentations for in-vivo MP2RAGE (left), postmortem MRI (middle) and 3D reconstructed histology. D, Cortical surface reconstructions (anterior-top; posterior-bottom) for in-vivo (top) and post-mortem data (bottom).

The segmentation algorithm further proved robust across a range of modalities, providing detailed cerebellar delineations not only in our own in-vivo 0.4mm isotropic MP2RAGE MRI, but also a post-mortem 0.2mm isotropic Fast Low Angle Shot MRI scan of the cerebellar cortex (n=1; seminal study reconstructing the cerebellar cortex; publicly available)^9^ and 3D-reconstructed (post-mortem) histology data (n=2; study combining multiple, coregistered, immunohistochemistry modalities; publicly available)^44^. While the higher-resolution 12h post-mortem MRI data allowed extracting visibly more cortical detail, our in-vivo 18.5min MRI scan still allowed delineating the cerebellar gray matter in detail, resolving many, though not all, individual folia. As a further visual validation, the cerebellar cortical surfaces of the in-vivo and post-mortem data were reconstructed (Fig.2D). In both in-vivo and post-mortem data, the cerebellar cortical surfaces retained the known characteristics from anatomical studies, including resolving the fissures which overlay the parallel fibers (giving the human cerebellum a characteristic canonically-ribbed appearance in the mediolateral direction), as well as splitting folia in the lateral side (black arrows, bottom Fig.2D).

These detailed segmentations enabled probing the regional cerebellar morphology and vascularization in-vivo and post-mortem. Importantly, we also leveraged the segmentation of the 3D-reconstructed (immuno)histology to compare different aspects of the cerebellar cytoarchitecture with the in-vivo measures (Fig.2A-C).

### The cerebellar cortical morphology is regionally heterogeneous

The combination of our in-vivo cerebellar imaging and segmentation allowed us to estimate the cerebellar cortical thickness (Fig.3A) and probe the regional cerebellar morphology. The cerebellar cortex showed a marked heterogeneity in its regional cortical thickness between lobules (Friedman test, X_2_(10)=21.9, p=0.016), with increased cortical thickness in lobules VIIIA (median [IQR] = 1.03 [0.87-1.13]), the medial side of Lobule VIIB (median [IQR] = 1.1 [0.95-1.23]), as well as local regions of Crus II (median [IQR] = 1.15 [0.93-1.2]; individual-level projection: Fig.3B, group median estimates: Fig.3C, Table E1). This result was reproduced in our segmentation of the post-mortem MRI data, with increased cortical thickness in VIIIA, VIIB and Crus II; Fig3B), despite a different MR contrast and resolution.

**Figure 3:**
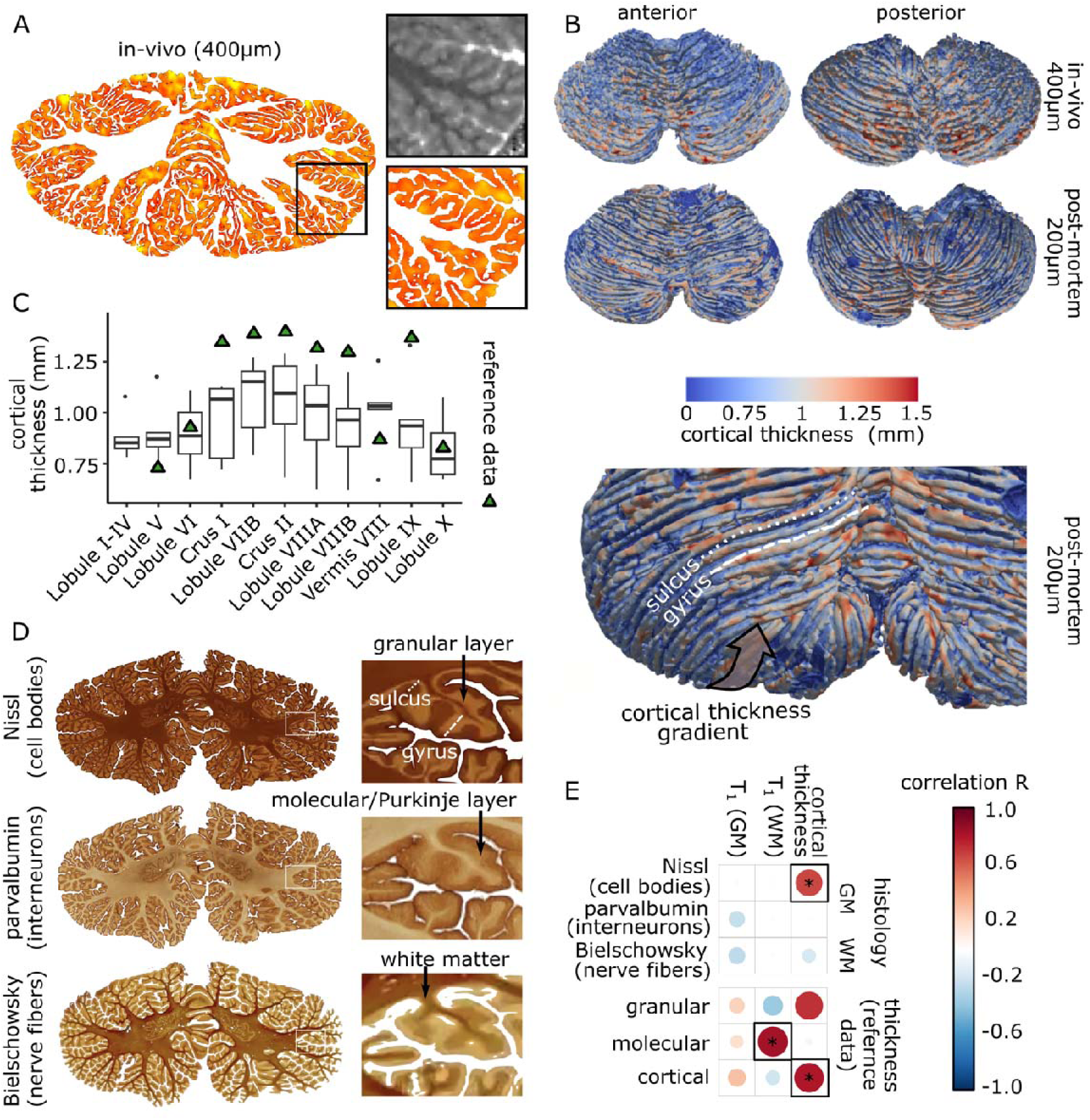
A, Cortical thickness estimate and comparison with MP2RAGE. B, Surface-projected cortical thickness in-vivo (top) and post-mortem (bottom). C, In-vivo median lobular thickness estimates versus reference data^8^ (triangles). D, The distinct immunohistology modalities allowed probing different aspects of the cerebellar cytoarchitecture. E, Correlations between in-vivo structural measures, immunohistology measures and reference data^8^. Significant correlations following FDR are encircled and marked with an asterisk *

The in-vivo group median of the lobular thickness significantly correlated with the lobular thickness derived from our segmentations of the 3D-reconstructed histology and post-mortem MRI data (S-Fig.1D-E), further suggesting that the derived morphological measures are consistent across modalities.

We validated our in-vivo estimate by comparing it with the only other whole-cerebellum morphology study known to us (n=1, 3D-reconstructed histology from the BigBrain^8,38^). Our median in-vivo estimate of the cerebellar cortical thickness strongly correlated with these reference data (R=0.81, p_FDR-corrected_=0.02, Fig.3F; triangles in Fig.3C), supporting that our in-vivo estimate of the cortical thickness truly reflects the underlying morphology. Our median estimate was slightly attenuated compared to the reference data by 0.25mm. This may reflect either increased partial volume effects from the necessarily lower resolution in in-vivo acquisition or better describe the in-vivo population (the reference data consist of n=1 histology). Together, our results suggest that there is regional heterogeneity in the cerebellar cortical morphology, which can be furthermore measured in-vivo with MRI (group-level results: S-Fig.2).

The cerebellar cortical regional morphology relates to the cerebellar cytoarchitecture We subsequently examined how the regional cerebellar morphology relates to the cerebellar cytoarchitecture. The (immuno)histological stainings we examined are sensitive to different aspects of the cerebellar lamination: the polarized (i.e. inverted) Nissl staining is particularly sensitive to the cell soma in the granular layer i.e. the granule and Golgi cells (increased intensity with higher cell body density), parvalbumin immunoreactivity labels the interneurons present in the molecular and Purkinje layers, while the Bielschowsky staining was used to labels axons in the white matter.

To correlate microscopy results with the in-vivo measures, we then extracted lobule-level average intensity estimates from the 3D-reconstructed (immuno)histology^44^ using the cerebellar segmentation. We found that the in-vivo cortical thickness strongly correlated with the lobular signal-intensity of the Nissl stainings (R=0.66, p_FDR-corrected_ <0.001, Table E2, Fig.3E), indicating a positive relationship between in-vivo cortical thickness and cell density of the granular layer.

The median T_1_ in white matter and gray matter also showed lobular-specific variability (S-Fig.2, Table E1). Furthermore, a positive relationship between T_1_ of the white matter and the reference molecular layer thickness was found, R=0.83, p_FDR-corrected_=0.02, Table E2, but no significant correlation to our post-mortem cytoarchitecture lobular measures was found.

### The regional cerebellar vascularization relates to the cerebellar cortical morphology and cerebellar cytoarchitecture

We then segmented the cerebellar arteries and veins by applying a recursive 2D ridge diffusion filter^46^ to the in-vivo TOF-MRA and venograms (Fig.4A-B, S-Fig.2, Table E1). The visualized cerebellar vasculature followed the reported anatomical description^47,48^ with the main arteries (Fig.4C), as well as superficial and deep veins being clearly visible. Our reconstruction of the cerebellar cortical surface and vascular system revealed that the cerebellar surface veins frequently run within the cerebellar fissures, i.e. between the cerebellar folia (Fig.4D-E), which are consistently mediolaterally oriented.

**Figure 4:**
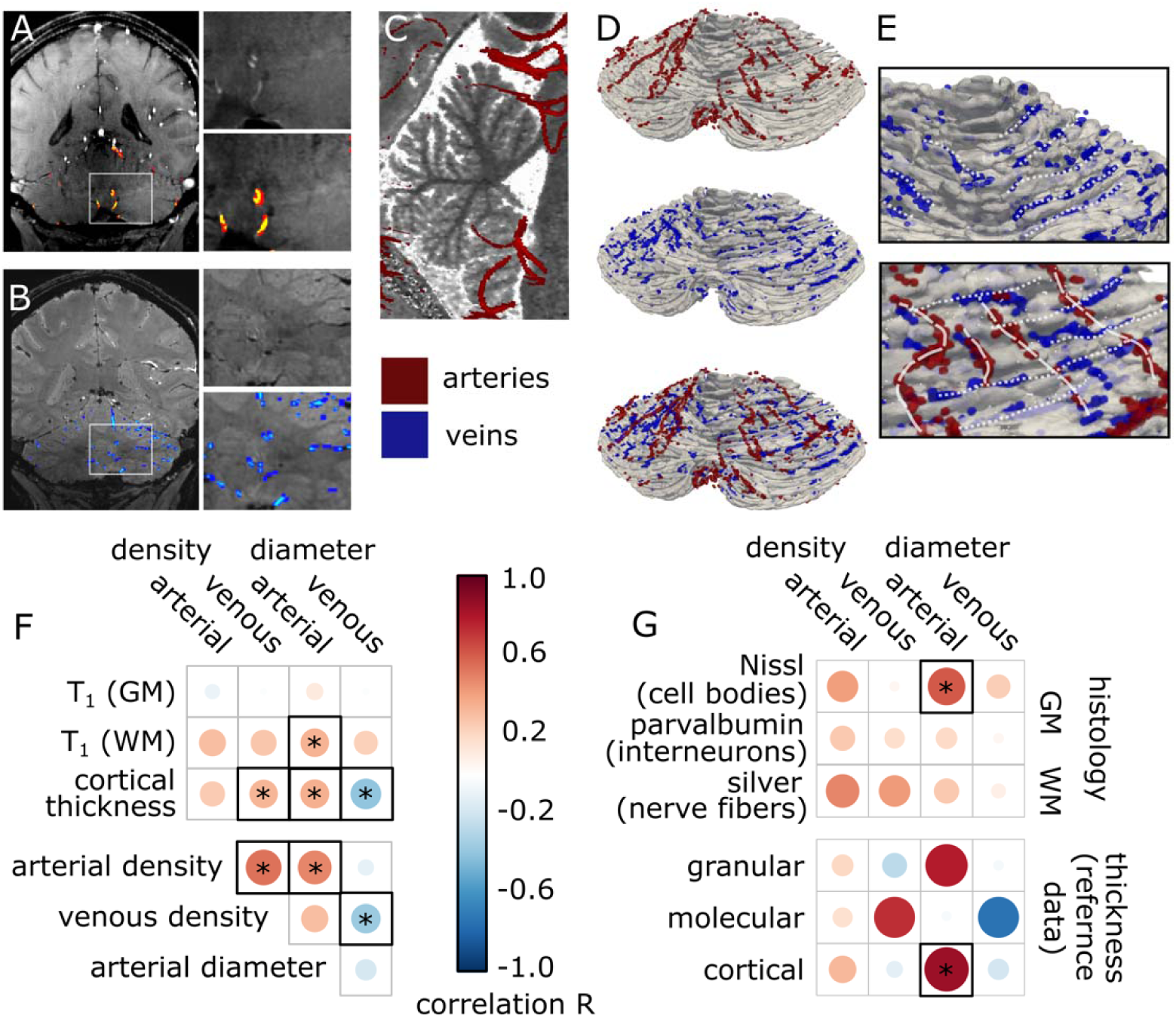
A, Time-of-flight and venogram (B) images and matching arterial (red-yellow) and venous (blue-light blue) segmentations. C, Maximum intensity projection of time-of-flight overlaid on T_1_ map. The feeding arteries of the cerebellum can be seen. D, Arterial (red) and venous (blue) 3D reconstructions and close-ups (E). F, Correlations between in-vivo structural-vascular measures. G, Correlations between in-vivo vascular measures, histology measures and reference data^8^. Statistically significant correlations following FDR are marked with *

Across the group, the lobular arterial density correlated with the lobular venous density (R=0.54, p_FDR-corrected_<0.001, Table E3, Fig.4F), in line with an expected relationship between arterial blood supply and venous blood drainage. This correlation estimate is identical to the venous-arterial density correlation reported for the neocortex (R=0.54)^20^, which supports the accuracy of our vascular tracing.

We then compared our reconstruction of the regional cerebellar vasculature to the cerebellar cortical morphology and cytoarchitecture. In our in-vivo data, we found positive correlations between vascularization density and cortex thickness (venous density-thickness: R=0.34, p_FDR-corrected_=0.04; arterial diameter-thickness: R=0.34, p_FDR-corrected_=0.04; Table E3, Fig.4F). This relationship between vascularization and morphology was confirmed when examining the cortical thickness reported from the BigBrain post-mortem data (arterial diameter-Big brain thickness: R=0.85, p_FDR-corrected_=0.02; a trend for arterial diameter-Big brain granular layer thickness: R=0.78, p_FDR-corrected_=0.05; Table E4, Fig.4F). This consistent relationship between vascularization and both in-vivo and post-mortem cortical morphology suggests that the regional vascularization of the cerebellar cortex is linked to morphology.

Probing further the relationship between cytoarchitecture and vascularization revealed a positive correlation between the in-vivo vascularization and signal-intensity of the Nissl staining (arterial diameter-Nissl staining intensity: R=0.61, p_FDR-corrected_=0.01; and a trend for arterial density-Nissl staining intensity: R=0.42, p_FDR-corrected_=0.08; Table E2, Fig.3E). This suggests that the observed relationship between vascularization and morphology may be driven by the functional needs or development of the granular layer, which is the main driver of Nissl stain intensity in the cerebellum.

We also detected a negative correlation between venous diameter and in-vivo cortical thickness (R=-0.4, p_FDR-corrected_=0.02), a trend for a positive correlation between arterial density and Bielschowsky-stain intensity (i.e. increased cortical arterial density correlates with reduced nerve-fiber density in white matter, R=0.49, p_FDR-corrected_=0.05) and a trend with BigBrain post-mortem molecular layer thickness (venous diameter-molecular layer thickness; R=-0.73, p_FDR-corrected_=0.05; venous density-molecular layer thickness: R=0.73, p_FDR-corrected_=0.05). This was also echoed in the in-vivo morphology (arterial diameter-T_1_wm: R=0.34, p_FDR-corrected_=0.04; and a trend for arterial density-T_1_wm: R=0.3, p_FDR-corrected_=0.06).

The vascular architecture of the brain shows high pial vascular density, while larger vessels tend to penetrate deeper. This produces a relative increase in vessel diameter and reduced vessel density deeper in the cortex^48^. We therefore purport that these correlations relate to this higher-vessel density and relative decrease in vessel diameter close to the cerebellar cortical surface (i.e. the molecular layer).

## Discussion

The cerebellum contains the majority of the human brain neurons, packed in a thin and tightly-folded cortex. This cortical area was largely out of reach of in-vivo imaging. For the first time, we comprehensively imaged the human cerebellar cortex by combining several key technologies of MR imaging and analysis. Our findings challenge the commonly-held assumption that the cerebellar cortex is uniform^5^, revealing instead a convergent interlobular heterogeneity in cortical morphology and vasculature.

Comparison of the in-vivo human morphology with post-mortem morphological and cytoarchitecture measures revealed that this heterogeneity likely arises from interlobular thickness and cell density variation in the granular layer. The granular layer consists of ∼50 billion granular cells, the most common neurons in the human brain^49^. While seemingly homogeneous, their density has been shown to vary regionally in several animal species^50,51^, in line with our human results.

The causal mechanism behind the correlation between the granular layer cell density and cerebellar cortical thickness is not clear. Similar relationships in the neocortex are thought to be mediated by the neuropil (e.g. axons and dendrites) which forms the largest part of the cortical volume rather than cell bodies^16–18^. In support of a similar interpretation, emerging evidence suggests that GPRIN3 gene expression, which is crucial for the development of neuron projection, exhibits interlobular variation^52^. This GPRIN3 transcriptomic gradient, which is conserved across mice and humans, seemingly aligns on a lobular level with the cortical thickness gradient we report^52^.

In the neocortex, morphological measures such as cortical thickness reflect not only the laminar architecture but also processing hierarchies, such as those established in sensory integration^53,54^. These hierarchies arise from computational specialization and variations in interregional connectivity^55,56^. Equivalent hierarchies are much less explored in the cerebellum, though functional gradients have been reported in humans using fMRI^57^.

The cortical heterogeneity we observe here mirrors these fMRI gradients, suggesting a link between the cortical morphology and cerebellar function. Recent evidence indicates significant variation in the convergence of neocortical inputs to the cerebellar cortex, which may drive the emergence of sensory integration in the cerebellum^58,59^. Future research may explore potential links between granular cell type, cortical morphology and function similarly to relationships between cerebellar transcriptomic and functional gradients reported in rats^7^.

The granular layer is a key determinant of cerebellar energy consumption, with 80% of cerebellar energy usage attributed to its signal transmission^60^. The granular layer therefore likely drives the cerebellar vascular demands^6^. Consistent with this, we found that cerebellar vascular density positively correlates with granular layer thickness and cytoarchitecture, establishing a link between the anatomical substrate and metabolic needs, similar to associations reported in the neocortex^20^ and hippocampus^21^. Such links evolve during development, during which the blood brain barrier and vascular branching is shaped by signals from the neural tissue^61^. These also guide the vessel alignment with the cerebellar folds, which had previously been reported post-mortem, and which we observed here in-vivo. Note that, based on the BOLD-fMRI contrast mechanism, the preference for a mediolateral orientation we observed in the cerebellar surface vessels will produce a similarly oriented bias in fMRI clusters. This is in addition to the known surface bias that is commonly observed in the cerebral cortex and is important to consider when interpreting high-resolution cerebellar fMRI results.

Importantly, our findings also demonstrate that our cortical thickness estimate, a measure we derived from data acquired in-vivo in a patient-relevant time frame (18min) and with largely automated acquisition and processing pipelines, is both faithful to the true cortical thickness and sensitive to the cerebellar granular layer cell density. By comparison, efforts to probe the cerebellar cytoarchitecture with histology or postmortem MRI required laborious manual work, resulting in a dearth of such data^8,9^. Crucially, it may be clinically important to probe the granular layer in-vivo: histopathological evidence suggests that in progressive multiple sclerosis, the granular layer experiences a high incidence of lesions and dramatic demyelination, which is more pronounced than in the neocortex^62^. Due to the challenges associated with cerebellar imaging, this pathology has remained largely inaccessible in-vivo. However, it is likely linked to both motor impairment and cognitive deficits in multiple sclerosis^63,64^. Cerebellar cortical pathology has also been reported in several other neurodegenerative diseases, ranging from cerebellar ataxias to Alzheimer’s disease^65,66^. The cerebellar imaging framework we developed may enable investigations into these pathologies, akin to the recent progress made in neocortical imaging of laminar-specific multiple sclerosis pathology^67^. Note that achieving full resolution of individual cerebellar folia remains beyond the current MRI sensitivity and further technical advancements, such as higher field strengths, are likely necessary to overcome these limitations^68^.

In this study, we spanned several scales, from post-mortem cell density to mesoscale patterns measurable with in-vivo MRI, such as cortical thickness and vascularization. Our results show a remarkable spatial convergence across these features in the human cerebellar cortex. Future work may address how these convergences occur and leverage them to interpret structural and functional measures in health and disease.

## Methods

MRI Acquisition. 10 participants (6 women; age range = 25-45 years, median = 31) were scanned in a 7T Achieva MRI scanner (Philips Healthcare, Best, the Netherlands) using an 8 channel transmit and 32 channel receive whole-head coil (Nova Medical, Wilmington, USA). All participants provided written informed consent. The participants were screened before the experiments to confirm MRI compatibility. Approval of the experimental protocol was obtained from the local ethics committee.

The cerebellar cortex was imaged with an MP2RAGE protocol with a nominal resolution of 0.4mm isotropic, similar to our previous study^30^ (FOV=210×120×60 mm^3^; matrix size=524×300×150; echo time=3.1ms; repetition time=7ms; inversion time 1/inversion time 2=1000/2900 msec; flip angle for inversion times 1 and 2 respectively=7°/5°; sensitivity encoding factor y/z=1.5/1; MP2RAGE shot=5sec; acquisition time_MP2RAGE_=14.5 minutes). This two-readout inversion-recovery technique reduces the effect of the 7T B_1_ inhomogeneities (particularly challenging in the cerebellum^35^) while providing good GM/WM contrast^31^.

In between the MP2RAGE shots, whole-head fat navigators were acquired (3D echo-planar imaging acquisitions with a fat-selective binomial excitation pulse; FOV=240×240×120mm^3^; matrix=120×120×60; voxel size=2mm isotropic; echo time=1.88ms; repetition time=5.65ms; flip angle=1°; sensitivity encoding factor_y/z_=4/2; acquisition time_volume_=550msec). A motion correction algorithm^33^ realigned the reconstructed fat navigators in real time using a 6 degrees of freedom rigid body registration and updated the FOV (wait time=0.9 second), resulting in a total acquisition time=18.5 minutes for the fat-navigator interleaved MP2RAGE.

To optimize the B_1_^+^ transmit field over the cerebellum, which is known to be inhomogeneous for typical whole-head 7T coils^35^, we added phase offsets in each transmit channel of our parallel-transmit head coil compared to the phase shims in our standard (pseudo) circularly polarized mode. Typically, such phase offsets are optimized individually, i.e. in every scan session. To ensure RF-shimming robustness across group and minimize the scan time, we instead used a group-calculated phase shim optimized for the cerebellum over 12 previously-acquired datasets^36^. Briefly, these datasets consisted of a DREAM^69^ B_1_^+^ map, acquired while all coils were transmitting (FOV=224×224×168 mm^3^, voxel size=3.5mm isotropic, echo time=3ms, repetition time =16ms, flip angle=7°, acquisition time=20sec) as well as a spoiled gradient echo while transmitting with each channel sequentially (FOV=224×224×168 mm^3^, voxel size=3.5mm isotropic, echo time=1.97ms, repetition time=8ms, flip angle=1.5°, acquisition time=2min). From these, the relative channel-specific transmit fields were estimated to minimise the cost function sd(B_1+cerebellum_)/mean(B_1+cerebellum_)^2^ using the MRCodeTool (Tesla Dynamic Coils, Netherlands) and the median phase offsets versus the quadrature mode were used.

We have demonstrated the efficacy of both the motion correction algorithm^30,33^ and the population-level RF-shim approach before^36^. To visually demonstrate their contribution, in one participant we derived an MP2RAGE with and without motion correction and with and without RF-shimming (Fig.1C-D).

To visualize the cerebellar vasculature, we acquired a single-slab TOF (FOV=210x120x60mm^3^, voxel size=0.5mm isotropic, echo time=3.1ms, repetition time=10ms, sensitivity encoding factor_y/z_=3/1.5, flip-angle=18°, acquisition time=3min)^70^. The TOF was empirically optimized (pilot experiments in 2 participants) with regards to the slab size and FOV to reduce fresh blood saturation and maximize contrast. A FOV parallel to but not covering the anterior part of the brainstem (i.e. not covering the basilar artery) was found to provide the optimal contrast. We then implemented a spoiled gradient-recalled echo (GRE) sequence to visualize the cerebellar veins, matching the field of view and placement and approximately matching the spatial resolution of the TOF (FOV=210x120x60mm^3^, echo time=10ms, repetition time=15ms, flip angle=12°, voxel size=0.6mm isotropic, sensitivity encoding factor_y/z_=2.5/1, acquisition time =5min). The parameters for the venous-weighted GRE sequence were selected based on a previous 7T-optimized protocol^71^, which we adapted to a shorter echo time and reduced flip angle to reduce sensitivity to the increased dynamic B_0_ inhomogeneity in the cerebellum, while still retaining venous contrast (pilot experiments in 2 participants). In both scans, the group-optimized RF shims were employed to optimize the RF-excitation pattern. To retain the tissue saturation in the TOF and since the reduced spatial resolution makes motion less relevant, we did not employ prospective motion correction in the TOF and GRE scans.

At the start of each scan session, a B_0_ map was acquired (FOV=224×224×224mm^3^, voxel-size=3.5mm isotropic, echo time=1.54ms, repetition time=4ms, flip angle=8°, acquisition time=20sec). The B_0_ field was homogenized within the brain with a second-order shim by minimizing the variance of the B_0_ distribution in a least-squares way using MRCodeTool v1.5.7 (Tesla Dynamic Coils, Zaltbommel, The Netherlands). A whole-head lower-resolution magnetization-prepared rapid gradient-echo sequence was also acquired to ensure coverage of the cerebellum in the following scans and facilitate registration during processing (FOV=200×220×180mm^3^, matrix size=220×244×200, voxel size=0.9mm isotropic, repetition time=150ms, echo time=3ms, inversion time=1300ms, flip angle= 7°, sensitivity encoding factor_y/z_=2.5/2; acquisition time=2min).

Cerebellar cortical segmentation. The individual (inversion time 1 and inversion time 2) magnitude and phase MP2RAGE images were denoised using a local-complex principal components analysis denoising algorithm^42^ (Nighres 1.4.0)^41^, before the UNI image and T_1_ map were derived^31^.

The segmentation algorithm was applied on the T_1_ maps using functions implemented in Nighres. First, an initial rough segmentation of the cerebellum was performed: rigid-body transforms were calculated from the MP2RAGE to the whole-head MPRAGE, the TOF and GRE images, as well as a diffeomorphic transform from the whole-head MPRAGE to the MNI152 using the Advanced Normalization Tools 2.1.1^72^. Using the SUIT atlas^40^ a rough initial outline of the cerebellum and masks of individual lobules were projected to the 0.4mm isotropic T_1_ map. To facilitate smoother segmentations and surface reconstructions, the T_1_ maps were subsampled to 0.2mm isotropic. To reduce computational load, the T_1_ maps were cropped around the cerebellar ROI.

Then, a finer intensity-based segmentation step was applied: first, a fuzzy c-means clustering was applied to capture the large white matter (WM), gray matter (GM) and cerebrospinal fluid (CSF) cerebellar regions. In practice, we found that GM values had a bigger spread of values compared to the other classes. This, combined with remaining B_1_^+^ inhomogeneities, necessitated clustering with 4 classes (i.e. 2 clusters for GM, 1 cluster for WM and 1 cluster for CSF). A recursive 2D ridge diffusion filter^46^ was applied separately to capture bright and dark thin structures. This helped recover the thin 2D structures of CSF and WM that the tissue classification using fuzzy c-means missed due to partial voluming. In T_1_ maps, the cerebellar dura has a similar intensity profile as the thin WM branches. To derive a dura prior, we first defined a distance function from the cerebellar outline. We then estimated the dura as a 2D dark ridge, located close to the cerebellar mask outline and oriented parallel to that surface (as opposed to WM branches, which are also 2D dark ridges but oriented orthogonally to the outline). To facilitate topology correction in the next step, a partial-volumed version of all 2D ridges was created by dilating the probability maps by 0.4mm. We then defined WM, GM and CSF probability masks as:

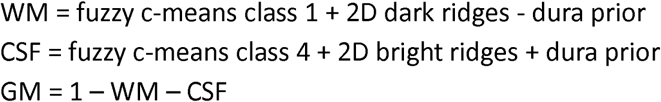

We finally performed topology correction of our segmentations: we first detected the largest connected object (WM), removing disconnected branches and holes. We used this to build a probability map of distance from the WM boundary using a sigmoid function. We combined this with the probability masks of the dilated WM, GM and CSF using an empirically-determined threshold of 50%. We then corrected the topology of the dilated WM probability map (i.e. spherical topology, non self-intersecting and smooth) using a fast-marching algorithm, starting from regions deeper inside the WM before handling the branches^73^. We finally found topology-correct surface boundaries of the target (non-dilated) WM, GM, and CSF probability maps using the CRUISE algorithm^43^. Briefly, CRUISE evolves a geometric deformable surface model starting from an initial topology-correct surface to the interfaces of cortical GM with WM and CSF, taking into account sulcal folding. As the initial topology-correct surface we used the topology-correct dilated WM probability map. A visual description of the segmentation process is described in S-Fig.1A. The segmentation scripts are shared in https://github.com/piloubazin/cerebellar-cortex.

The final topology-corrected WM and GM classifications were used to estimate the cerebellar cortical thickness, defined as the distance between the WM and GM surfaces. The mid-GM surface was tessellated into a dense triangulated mesh using a connectivity-consistent marching cube algorithm.

### Segmentation evaluation

The above segmentation pipeline was applied on all 10 in-vivo 0.4mm isotropic MP2RAGE datasets, providing consistent results as ascertained with visual checks. Note that there are no equivalent automated cerebellar segmentation approaches that aim to derive fine cerebellar details. For example, the current most popular cerebellar segmentation approach is optimized for ∼1mm datasets and is sensitive up to the large WM branches. As such, it is not an appropriate comparison. To evaluate the segmentation performance we therefore compared our segmentation’s output with a semi-manual, signal intensity-based approach without topology correction we had employed before^30^. The derived WM segmentations from the current pipeline were visible less noisy and the cortical surfaces visibly smoother (S-Fig.1B). We further applied our algorithm on the one post-mortem MRI dataset publicly available with a high-fidelity cerebellar surface reconstruction^9^. Compared to that segmentation (which was optimized for a smaller number of datasets and included laborious manual editing) our segmentation provided similar though qualitatively slightly worse results (i.e. less smooth, Fig.2C-D compared to Fig.2-3 in^9^). To check the generalizability of our approach to typical acquisitions, we also applied our segmentation approach to two whole head 7T MP2RAGE datasets available (0.64mm isotropic and 0.8mm isotropic, the last without RF shimming; S-Fig.2). Finally, to validate the derived cortical thickness estimates, we calculated the median across group per lobule and compared these with the one available quantification of regional cerebellar cortical thickness (based on N=1, 3D reconstructed histology from BigBrain^8^) using a Spearman correlation test. Statistical significance difference was established at p<0.05. To check if the cortical thickness estimate differed between lobules, the Friedman test was used, followed by pairwise Wilcoxon signed rank test between lobules. The results were corrected for multiple comparisons using the False Discovery Rate correction. All statistical analyses were carried out in R (v4.4.2).

Vascular imaging. From the TOF and GRE datasets, the cerebellar veins and arteries were delineated. This was achieved using a recursive 2D ridge diffusion filter (which enhances continuous vessels but penalizes inconsistent ones)^46^. The filter was applied to capture bright (arteries in TOF) and dark (veins in GRE) structures respectively. Using this filter, we extracted the arterial and venous vascular density per lobule ( # of arterial or venous voxels in lobule / # of voxels in lobule) and the average vessel diameter (estimated by combining the vessel segmentation with a Gaussian point spread function fit to account for partial-volume^46^) per lobule. From the MP2RAGE datasets, the lobular T_1_ values for the WM and GM ROIs were also extracted, besides the cortical thickness. The lobular morphological (T_1_ values, cortical thickness) and vascular (arterial and venous density and diameter) measures were compared using bootstrapped (n=5000) repeated-measures correlations (rmcorr package). The results were corrected for multiple comparisons using the False Discovery Rate correction.

In-vivo imaging and immunohistology comparison. To compare our in-vivo results with the cerebellar cytoarchitecture, we employed a dataset of 3D-reconstructed (immuno)histology, including Nissl, parvalbumin and Bielschowsky, previously published by the authors (technical details here: ^44^). This dataset included n=2 brains from non-demented donors. The segmentation algorithm was applied to the 3D-reconstructed Nissl image, producing delineations of the GM and WM, coregistered to all stainings^44^. To probe the regional cytoarchitectural properties, the lobular signal intensity was intensity was extracted. Given that Nissl stains for the cell bodies in the granular layer, parvalbumin labels the interneurons in the molecular/Purkinje layer and Bielschowsky is sensitive to the nerve fibers in the WM, the signal from the appropriate lobular ROIs was used (i.e. Nissl and parvalbumin: GM ROI, Bielschowsky: WM ROI). To increase tissue contrast in the Nissl sections, the images were acquired with a polarization filter, resulting in inverted intensity. To normalize the lobular signal intensity between the n=2 datasets, first a z-stat transform was applied to the intensity within dataset and then the median of the two datasets was calculated.

These normalized signal values from the (immuno)histology were subsequently compared with the in-vivo results, as well as the granular, molecular and total cortical thickness estimates from the BigBrain reference data^8^. To achieve that, first, the group median for each in-vivo morphological and vascular lobular measure was calculated. Then, Spearman correlations were performed these in-vivo medians and the normalized signal values from histology and BigBrain layer/cortical thickness estimates. The False Discovery Rate correction was applied for multiple comparisons.

## Acknowledgements

This work was supported by the NWO XS (OCENW.XS22.4.007) and Amsterdam Brain and Cognition Talent (T0922) grants to N.P.; and NWO TTW VIDI (VI.Vidi.198.016) and Aspasia (015.014.038) grants to W.v.d.Z.;

## References

1. Barton, R. A. & Venditti, C. Rapid Evolution of the Cerebellum in Humans and Other Great Apes. Curr. Biol. 24, 2440–2444 (2014).

2. Magielse, N. et al. Phylogenetic comparative analysis of the cerebello-cerebral system in 34 species highlights primate-general expansion of cerebellar crura I-II. Commun. Biol. 6, 1–17 (2023).

3. Azevedo, F. A. C. et al. Equal numbers of neuronal and nonneuronal cells make the human brain an isometrically scaled-up primate brain. J. Comp. Neurol. 513, 532–541 (2009).

4. Schmahmann, J. D. From movement to thought: Anatomic substrates of the cerebellar contribution to cognitive processing. Hum. Brain Mapp. 4, 174–198 (1996).

5. Voogd, J. & Glickstein, M. The anatomy of the cerebellum. Trends Cogn. Sci. 2, 307–313 (1998).

6. Diedrichsen, J., King, M., Hernandez-Castillo, C., Sereno, M. & Ivry, R. B. Universal Transform or Multiple Functionality? Understanding the Contribution of the Human Cerebellum across Task Domains. Neuron 102, 918–928 (2019).

7. Hao, S. et al. Cross-species single-cell spatial transcriptomic atlases of the cerebellar cortex. Science 385, eado3927 (2024).

8. Zheng, J. et al. Three-Dimensional Digital Reconstruction of the Cerebellar Cortex: Lobule Thickness, Surface Area Measurements, and Layer Architecture. The Cerebellum 22, 249–260 (2023).

9. Sereno, M. I. et al. The human cerebellum has almost 80% of the surface area of the neocortex. Proc. Natl. Acad. Sci. 117, 19538–19543 (2020).

10. Wolfs, E. M. L., Van der Zwaag, W., Priovoulos, N., Klaus, J. & Schutter, D. J. L. G. The cerebellum during provocation and aggressive behaviour: A 7 T fMRI study. Imaging Neurosci. 1, 1–18 (2023).

11. King, M., Hernandez-Castillo, C. R., Poldrack, R. A., Ivry, R. B. & Diedrichsen, J. Functional boundaries in the human cerebellum revealed by a multi-domain task battery. Nat. Neurosci. 22, 1371–1378 (2019).

12. Brouwer, E. J. P., Priovoulos, N., Hashimoto, J. & van der Zwaag, W. Proprioceptive engagement of the human cerebellum studied with 7T-fMRI. Imaging Neurosci. 2, 1–12 (2024).

13. Brodmann, K. Vergleichende Lokalisationslehre Der Grosshirnrinde in Ihren Prinzipien Dargestellt Auf Grund Des Zellenbaues. (Barth, 1909).

14. Broca, P. Remarques sur le siège de la faculté du langage articulé, suivies d’une observation d’aphémie (perte de la parole). Bull. Memoires Soc. Anat. Paris 6, 330–357 (1861).

15. Fischl, B. FreeSurfer. NeuroImage 62, 774–781 (2012).

16. Wagstyl, K. et al. Mapping Cortical Laminar Structure in the 3D BigBrain. Cereb. Cortex 28, 2551–2562 (2018).

17. la Fougère, C. et al. Where in-vivo imaging meets cytoarchitectonics: The relationship between cortical thickness and neuronal density measured with high-resolution [18F]flumazenil-PET. NeuroImage 56, 951–960 (2011).

18. Gong, G., He, Y., Chen, Z. J. & Evans, A. C. Convergence and divergence of thickness correlations with diffusion connections across the human cerebral cortex. NeuroImage 59, 1239–1248 (2012).

19. Tsai, P. S. et al. Correlations of Neuronal and Microvascular Densities in Murine Cortex Revealed by Direct Counting and Colocalization of Nuclei and Vessels. J. Neurosci. 29, 14553–14570 (2009).

20. Bernier, M., Cunnane, S. C. & Whittingstall, K. The morphology of the human cerebrovascular system. Hum. Brain Mapp. 39, 4962 (2018).

21. Haast, R. A. M. et al. Insights into hippocampal perfusion using high-resolution, multi-modal 7T MRI. Proc. Natl. Acad. Sci. 121, e2310044121 (2024).

22. Geschwind, D. H. & Rakic, P. Cortical Evolution: Judge the Brain by Its Cover. Neuron 80, 633–647 (2013).

23. Panizzon, M. S. et al. Distinct Genetic Influences on Cortical Surface Area and Cortical Thickness. Cereb. Cortex 19, 2728–2735 (2009).

24. Fjell, A. M. et al. Development and aging of cortical thickness correspond to genetic organization patterns. Proc. Natl. Acad. Sci. 112, 15462–15467 (2015).

25. McGinnis, S. M., Brickhouse, M., Pascual, B. & Dickerson, B. C. Age-Related Changes in the Thickness of Cortical Zones in Humans. Brain Topogr. 24, 279–291 (2011).

26. Walhovd, K. B., Fjell, A. M., Giedd, J., Dale, A. M. & Brown, T. T. Through Thick and Thin: a Need to Reconcile Contradictory Results on Trajectories in Human Cortical Development. Cereb. Cortex 27, bhv301 (2017).

27. Burgaleta, M., Johnson, W., Waber, D. P., Colom, R. & Karama, S. Cognitive ability changes and dynamics of cortical thickness development in healthy children and adolescents. NeuroImage 84, 810–819 (2014).

28. McDonald, C. R. et al. Regional neocortical thinning in mesial temporal lobe epilepsy. Epilepsia 49, 794–803 (2008).

29. Antel, S. B. et al. Automated detection of focal cortical dysplasia lesions using computational models of their MRI characteristics and texture analysis. NeuroImage 19, 1748–1759 (2003).

30. Priovoulos, N., Andersen, M., Dumoulin, S. O., Boer, V. O. & van der Zwaag, W. High-Resolution Motion-corrected 7.0-T MRI to Derive Morphologic Measures from the Human Cerebellum in Vivo. Radiology 307, e220989 (2023).

31. Marques, J. P. et al. MP2RAGE, a self bias-field corrected sequence for improved segmentation and T1-mapping at high field. NeuroImage 49, 1271–1281 (2010).

32. Bazin, P.-L. et al. Sharpness in motion corrected quantitative imaging at 7T. NeuroImage 222, 117227 (2020).

33. Andersen, M., Laustsen, M. & Boer, V. Accuracy investigations for volumetric head-motion navigators with and without EPI at 7 T. Magn. Reson. Med. 88, 1198–1211 (2022).

34. Raimondo, L. et al. Robust high spatio-temporal line-scanning fMRI in humans at 7T using multi-echo readouts, denoising and prospective motion correction. J. Neurosci. Methods 384, 109746 (2023).

35. Priovoulos, N. et al. A local multi-transmit coil combined with a high-density receive array for cerebellar fMRI at 7 T. NMR Biomed. 34, e4586 (2021).

36. Brouwer, E. J. P., Priovoulos, N. & Van der Zwaag, W. A universal B1 shim for the human cerebellum. in Proceedings of the Annual Meeting of ISMRM, Toronto, Canada, 2707 (2023).

37. Priovoulos, N. & Bazin, P.-L. Methods for cerebellar imaging analysis. Curr. Opin. Behav. Sci. 54, 101328 (2023).

38. Amunts, K. et al. BigBrain: An Ultrahigh-Resolution 3D Human Brain Model. Science 340, 1472–1475 (2013).

39. Edlow, B. L. et al. 7 Tesla MRI of the ex vivo human brain at 100 micron resolution. Sci. Data 6, 244 (2019).

40. Diedrichsen, J., Balsters, J. H., Flavell, J., Cussans, E. & Ramnani, N. A probabilistic MR atlas of the human cerebellum. NeuroImage 46, 39–46 (2009).

41. Huntenburg, J. M., Steele, C. J. & Bazin, P.-L. Nighres: processing tools for high-resolution neuroimaging. GigaScience 7, giy082 (2018).

42. Bazin, P.-L. et al. Denoising High-Field Multi-Dimensional MRI With Local Complex PCA. Front. Neurosci. 13, (2019).

43. Han, X. et al. CRUISE: Cortical reconstruction using implicit surface evolution. NeuroImage 23, 997–1012 (2004).

44. Alkemade, A. et al. A unified 3D map of microscopic architecture and MRI of the human brain. Sci. Adv. 8, eabj7892 (2022).

45. Zheng, J. et al. Three-Dimensional Digital Reconstruction of the Cerebellar Cortex: Lobule Thickness, Surface Area Measurements, and Layer Architecture. The Cerebellum 22, 249–260 (2023).

46. Bazin, P.-L., Plessis, V., Fan, A. P., Villringer, A. & Gauthier, C. J. Vessel segmentation from quantitative susceptibility maps for local oxygenation venography. in 2016 IEEE 13th International Symposium on Biomedical Imaging (ISBI) 1135–1138 (2016). doi:10.1109/ISBI.2016.7493466.

47. Delion, M., Dinomais, M. & Mercier, P. Arteries and Veins of the Cerebellum. The Cerebellum 16, 880–912 (2017).

48. Duvernoy, H., Delon, S. & Vannson, J. L. The vascularization of the human cerebellar cortex. Brain Res. Bull. 11, 419–480 (1983).

49. Consalez, G. G., Goldowitz, D., Casoni, F. & Hawkes, R. Origins, Development, and Compartmentation of the Granule Cells of the Cerebellum. Front. Neural Circuits 14, 611841 (2021).

50. Cerminara, N. L., Lang, E. J., Sillitoe, R. V. & Apps, R. Redefining the cerebellar cortex as an assembly of non-uniform Purkinje cell microcircuits. Nat. Rev. Neurosci. 16, 79–93 (2015).

51. Lange, W. Regional differences in the distribution of golgi cells in the cerebellar cortex of man and some other mammals. Cell Tissue Res. 153, 219–226 (1974).

52. Kozareva, V. et al. A transcriptomic atlas of mouse cerebellar cortex comprehensively defines cell types. Nature 598, 214–219 (2021).

53. Wagstyl, K., Ronan, L., Goodyer, I. M. & Fletcher, P. C. Cortical thickness gradients in structural hierarchies. NeuroImage 111, 241–250 (2015).

54. Grill-Spector, K. & Malach, R. THE HUMAN VISUAL CORTEX. Annu. Rev. Neurosci. 27, 649–677 (2004).

55. Barbas, H. Pattern in the laminar origin of corticocortical connections. J. Comp. Neurol. 252, 415–422 (1986).

56. Barbas, H. & Rempel-Clower, N. Cortical structure predicts the pattern of corticocortical connections. Cereb. Cortex 7, 635–646 (1997).

57. Guell, X., Schmahmann, J. D., Gabrieli, J. D. & Ghosh, S. S. Functional gradients of the cerebellum. eLife 7, e36652 (2018).

58. King, M., Shahshahani, L., Ivry, R. B. & Diedrichsen, J. A task-general connectivity model reveals variation in convergence of cortical inputs to functional regions of the cerebellum. eLife 12, e81511 (2023).

59. Lindeman, S., et al. Cerebellar Purkinje cells can differentially modulate coherence between sensory and motor cortex depending on region and behavior. Proc. Natl. Acad. Sci. 118, e2015292118 (2021).

60. Howarth, C., Peppiatt-Wildman, C. M. & Attwell, D. The Energy Use Associated with Neural Computation in the Cerebellum. J. Cereb. Blood Flow Metab. 30, 403–414 (2010).

61. Liebner, S. & Plate, K. H. Differentiation of the brain vasculature: the answer came blowing by the Wnt. J. Angiogenesis Res. 2, 1 (2010).

62. Kutzelnigg, A. et al. Widespread Demyelination in the Cerebellar Cortex in Multiple Sclerosis. Brain Pathol. 17, 38–44 (2007).

63. Parmar, K. et al. The role of the cerebellum in multiple sclerosis—150 years after Charcot. Neurosci. Biobehav. Rev. 89, 85–98 (2018).

64. Brouwer, E. J., Strik, M. & Schoonheim, M. M. The role of the cerebellum in multiple sclerosis: structural damage and disconnecting networks. Curr. Opin. Behav. Sci. 59, 101436 (2024).

65. Pagen, L. H. G. et al. Contributions of Cerebro-Cerebellar Default Mode Connectivity Patterns to Memory Performance in Mild Cognitive Impairment. J. Alzheimers Dis. 75, 633–647 (2020).

66. Koeppen, A. H. The pathogenesis of spinocerebellar ataxia. The Cerebellum 4, 62–73 (2005).

67. Mainero, C. et al. A gradient in cortical pathology in multiple sclerosis by in vivo quantitative 7 T imaging. Brain 138, 932–945 (2015).

68. Bates, S. et al. A vision of 14 T MR for fundamental and clinical science. Magn. Reson. Mater. Phys. Biol. Med. 36, 211–225 (2023).

69. Nehrke, K., Versluis, M. J., Webb, A. & Börnert, P. Volumetric B1 (+) mapping of the brain at 7T using DREAM. Magn Reson Med 71, 246–256 (2014).

70. de Buck, M. H. S., Jezzard, P. & Hess, A. T. Optimization of undersampling parameters for 3D intracranial compressed sensing MR angiography at 7 T. Magn. Reson. Med. 88, 880– 889 (2022).

71. Koopmans, P. J., Manniesing, R., Niessen, W. J., Viergever, M. A. & Barth, M. MR venography of the human brain using susceptibility weighted imaging at very high field strength. Magma N. Y. N 21, 149–158 (2008).

72. Avants, B. B. et al. A reproducible evaluation of ANTs similarity metric performance in brain image registration. NeuroImage 54, 2033–2044 (2011).

73. Bazin, P.-L. & Pham, D. L. Topology-Preserving Tissue Classification of Magnetic Resonance Brain Images. IEEE Trans. Med. Imaging 26, 487–496 (2007).

