## Supplemental Material for "Mesoscale imaging of the human cerebellum reveals converging regional specialization of its morphology, vasculature and cytoarchitecture"

Supplementary material

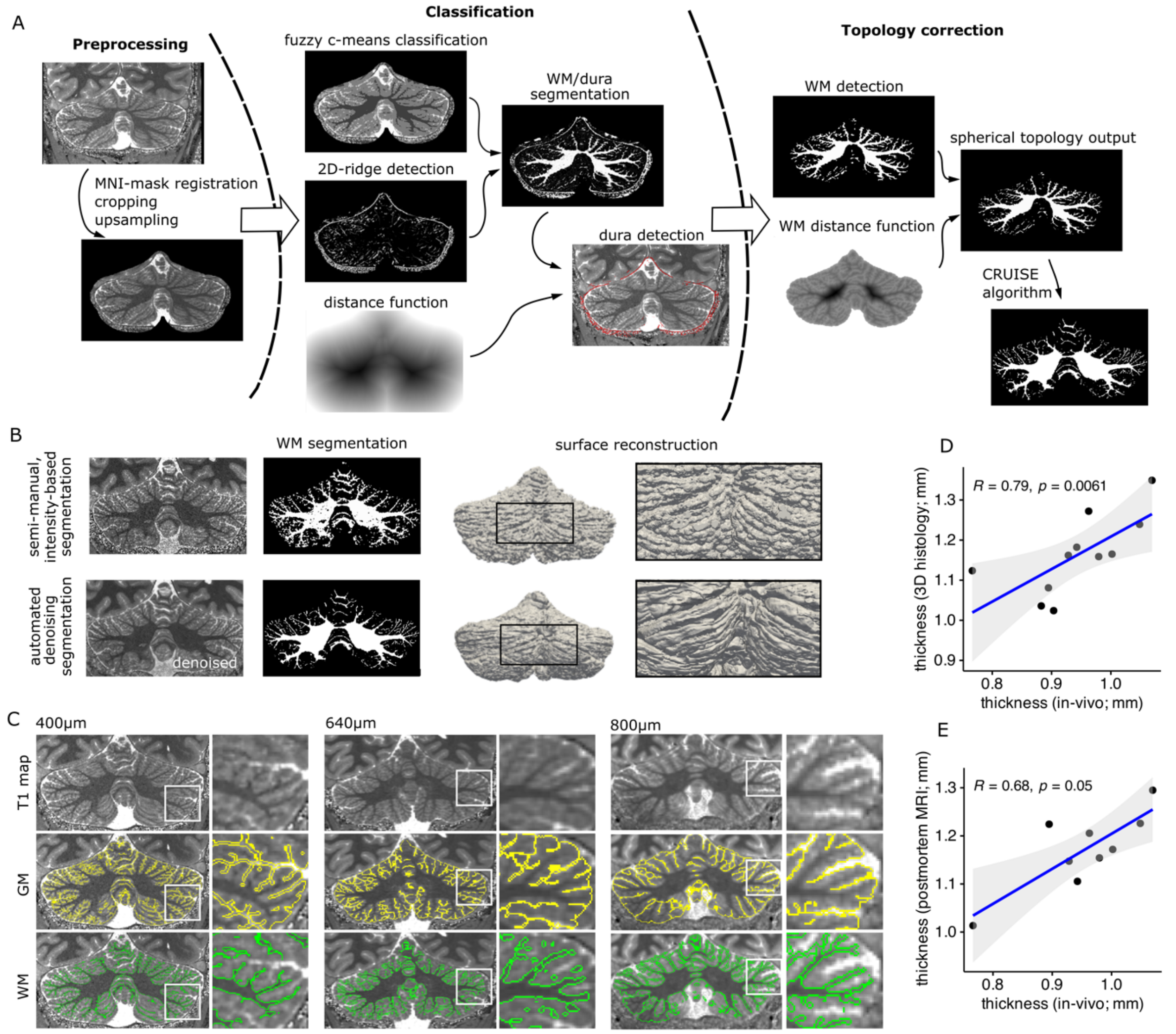


*S-Figure 1: A, Segmentation pipeline. B, Segmentation comparison between previous semi-manual segmentations[3] and our automated pipeline. C, Comparison of segmentation results between our 0.4mm isotropic resolution MP2RAGE and typical whole-head MP2RAGE protocols at 7T at 0.64mm and 0.8mm without RF phase-shimming. D, Scatterplot and Spearman correlation between group median lobular thickness in in-vivo group and lobular thickness from the segmentation of postmortem MRI data and E, lobular thickness from our segmentation of 3D-reconstructed histology data*

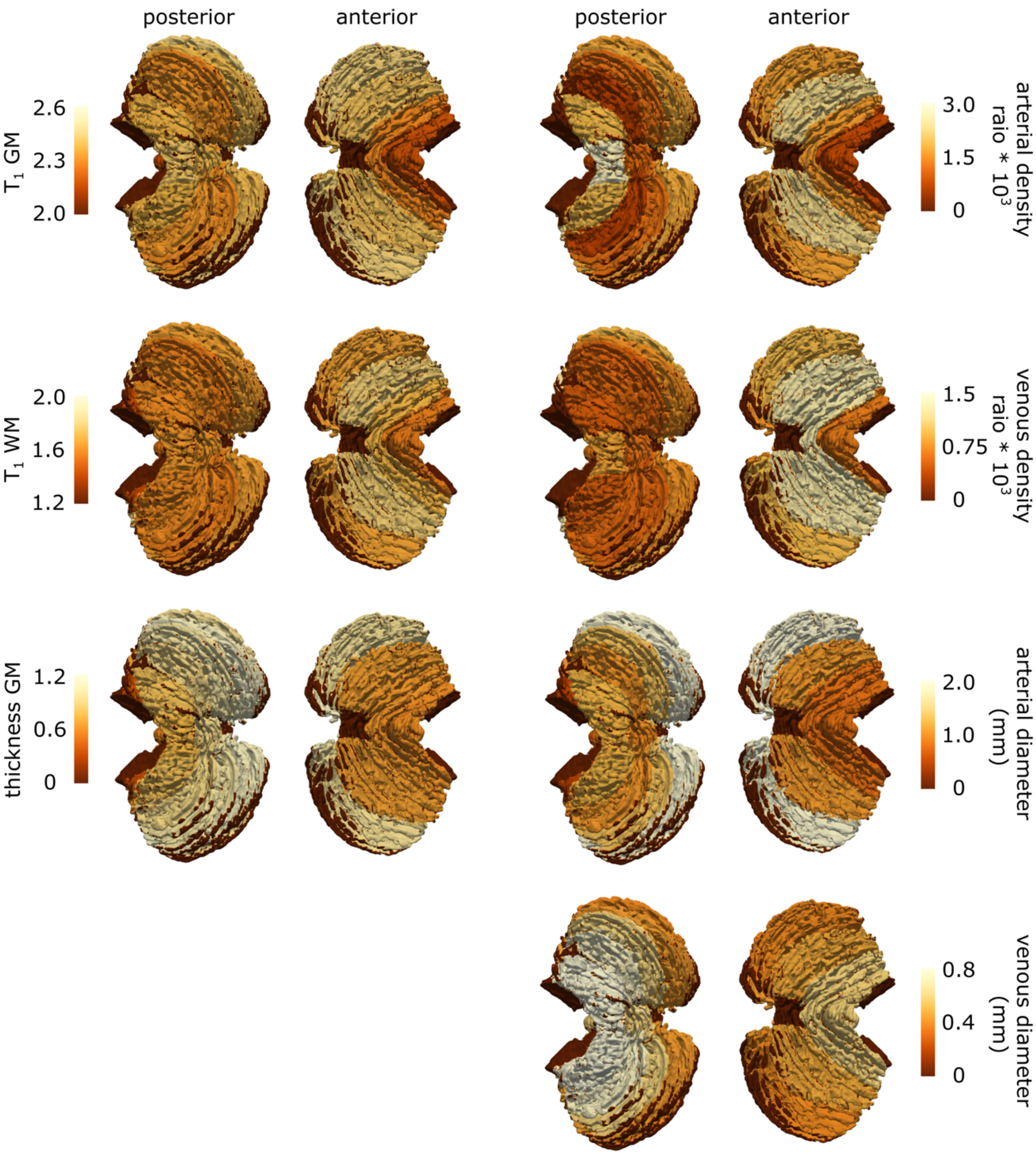


*S-Figure 2: Median estimates of in-vivo morphological and vascular measures per lobule.*

| ROI | T1 (s) | | Volume ($cm^3$) | | | Thickness (cm) | Density (ratio *$10^3$) | | Diameter (mm) | |
|---|---|---|---|---|---|---|---|---|---|---|
| | GM | WM | Total | GM | WM | GM | Arteries | Veins | Arteries | Veins |
| Lobule I - IV | 2.18 (1.94-2.31) | 1.56 (1.47-1.65) | 4.73 (4.44-5.04) | 2.77 (2.37-2.98) | 1.61 (1.46-2.55) | 0.85 (0.82-0.88) | 0.83 (0.65-1.1) | 6.73 (5.8-9.47) | 0.81 (0.79-1.01) | 0.64 (0.43-0.64) |
| Lobule V | 2.29 (1.93-2.37) | 1.7 (1.5-1.75) | 4.97 (4.88-6.05) | 3.53 (3.51-3.96) | 1.63 (1.23-1.78) | 0.87 (0.83-0.9) | 1.62 (1.49-2.35) | 13.11 (12.01-14.28) | 0.81 (0.66-0.97) | 0.64 (0.46-0.66) |
| Lobule VI | 2.4 (2.02-2.47) | 1.78 (1.48-1.82) | 11.7 (11.24-13.89) | 9 (8.2-9.13) | 3.25 (3.04-3.93) | 0.89 (0.8-1) | 2.34 (1.54-2.47) | 13 (12.1-14.84) | 1.08 (0.9-1.72) | 0.55 (0.55-0.65) |
| Crus I | 2.44 (2.18-2.54) | 1.65 (1.55-1.83) | 18.5 (18.13-18.77) | 13.22 (13.16-14.34) | 5.28 (4.98-5.8) | 1.07 (0.78-1.12) | 1.67 (1.37-2.14) | 9.53 (9.32-9.56) | 1.96 (1.87-2.88) | 0.49 (0.49-0.59) |
| Lobule VIIB | 2.3 (2.14-2.41) | 1.56 (1.51-1.67) | 7.69 (6.59-7.88) | 5.05 (4.61-6.11) | 1.98 (1.85-1.98) | 1.1 (0.95-1.23) | 1.23 (0.24-1.53) | 5.6 (5.08-6.59) | 1.34 (0.93-1.61) | 0.64 (0.58-0.68) |
| CrusI I | 2.39 (2.18-2.47) | 1.64 (1.51-1.75) | 14.18 (13.33-15.82) | 10.09 (9.92-12.29) | 4.08 (3.53-4.21) | 1.15 (0.93-1.2) | 1.76 (1.17-2.26) | 7.7 (6.14-7.71) | 1.96 (1.78-2.08) | 0.52 (0.48-0.58) |
| Lobule VIIIA | 2.32 (2.15-2.41) | 1.59 (1.44-1.64) | 7 (6.23-7.69) | 4.53 (4.4-5.2) | 1.95 (1.82-2.44) | 1.03 (0.87-1.13) | 0.69 (0.67-2.39) | 5.85 (4.27-6.75) | 1.06 (1.01-1.43) | 0.76 (0.73-0.87) |
| Lobule VIIIB | 2.37 (2.09-2.44) | 1.6 (1.42-1.7) | 6.31 (5.28-6.41) | 3.71 (3.62-3.94) | 2.46 (1.66-2.58) | 0.97 (0.84-1.02) | 1.9 (0.82-2.04) | 5.16 (4.67-5.27) | 1.25 (1.21-1.47) | 0.89 (0.76-0.93) |
| Vermis VIII | 2.38 (2.04-2.47) | 1.63 (1.46-1.76) | 1.29 (1.2-1.35) | 0.9 (0.9-0.95) | 0.39 (0.31-0.4) | 1.03 (1.02-1.05) | 1.17 (0.51-2.19) | 6.39 (4.84-8.68) | 0.98 (0.95-1) | 0.64 (0.63-0.83) |
| Lobule IX | 2.41 (2.04-2.46) | 1.63 (1.43-1.71) | 4.54 (3.99-4.95) | 2.94 (2.91-2.98) | 1.6 (1.11-1.97) | 0.94 (0.83-0.97) | 2.76 (2.38-3.16) | 8.53 (7.37-9.68) | 1.21 (1.17-1.28) | 0.83 (0.79-0.9) |
| Lobule X | 2.37 (2.16-2.4) | 1.42 (1.39-1.56) | 0.78 (0.69-0.82) | 0.24 (0.17-0.3) | 0.51 (0.46-0.55) | 0.71 (0.67-0.84) | 0.15 (0.01-0.21) | 0.39 (0.14-0.57) | 0.51 (0.45-0.54) | 0.97 (0.73-1.11) |

*Table E1: Morphological and vascular characteristics of the cerebellar lobules*

| Measure 1 (post-mortem) | | Measure 2 (in-vivo) | correlation coefficient | p-value FDR adjusted |
|---|---|---|---|---|
| Reference thickness | granular | T1 GM | 0.22 | 0.74 |
| | molecular | T1 GM | 0.16 | 0.74 |
| | cortical | T1 GM | 0.3 | 0.73 |
| | granular | T1 WM | -0.37 | 0.66 |
| | molecular | T1 WM | 0.83 | **0.02** |
| | cortical | T1 WM | -0.2 | 0.74 |
| | granular | thickness GM | 0.72 | 0.06 |
| | molecular | thickness GM | 0.05 | 0.89 |
| | cortical | thickness GM | 0.81 | **0.02** |
| histology | Parvalbumin GM | T1 GM | -0.24 | 0.66 |
| | Nissl GM | T1 GM | -0.03 | 0.95 |
| | Silver WM | T1 GM | -0.26 | 0.66 |
| | Parvalbumin GM | T1 WM | 0.02 | 0.95 |
| | Nissl GM | T1 WM | -0.01 | 0.95 |
| | Silver WM | T1 WM | 0.02 | 0.95 |
| | Parvalbumin GM | thickness GM | 0.02 | 0.95 |
| | Nissl GM | thickness GM | 0.66 | **<0.001** |
| | Silver WM | thickness GM | -0.18 | 0.82 |

*Table E2: Spearman correlations between in-vivo morphological cerebellar measures and post-mortem cytoarchitecture and reference morphological measures.*

| Measure 1 | Measure 2 | correlation | p-value FDR |
|---|---|---|---|
| T1 GM | arterial density | -0.09 | 0.61 |
| T1 WM | arterial density | 0.3 | 0.06 |
| thickness GM | arterial density | 0.26 | 0.11 |
| T1 GM | venous density | -0.01 | 0.92 |
| T1 WM | venous density | 0.28 | 0.09 |
| thickness GM | venous density | 0.34 | **0.04** |
| arterial density | venous density | 0.54 | **<0.001** |
| T1 GM | arterial diameter | 0.11 | 0.55 |
| T1 WM | arterial diameter | 0.34 | **0.04** |
| thickness GM | arterial diameter | 0.35 | **0.04** |
| arterial density | arterial diameter | 0.49 | **<0.001** |
| venous density | arterial diameter | 0.31 | 0.06 |
| T1 GM | venous diameter | -0.02 | 0.92 |
| T1 WM | venous diameter | 0.23 | 0.15 |
| thickness GM | venous diameter | -0.4 | **0.02** |
| arterial density | venous diameter | -0.1 | 0.57 |
| venous density | venous diameter | -0.36 | **0.04** |
| arterial diameter | venous diameter | -0.18 | 0.29 |

*Table E3: Repeated-measures correlations between in-vivo vascular and in-vivo morphological cerebellar measures*

| Measure 1 (post-mortem) | | Measure 2 (in-vivo; vascular) | correlation coefficient | p-value FDR adjusted |
|---|---|---|---|---|
| Reference thickness | granular | arterial density | 0.2 | 0.85 |
| | molecular | arterial density | 0.17 | 0.85 |
| | cortical | arterial density | 0.32 | 0.85 |
| | granular | venous density | -0.26 | 0.85 |
| | molecular | venous density | 0.73 | 0.05 |
| | cortical | venous density | -0.12 | 0.9 |
| | granular | arterial diameter | 0.78 | 0.05 |
| | molecular | arterial diameter | -0.04 | 0.92 |
| | cortical | arterial diameter | 0.85 | **0.02** |
| | granular | venous diameter | -0.04 | 0.92 |
| | molecular | venous diameter | -0.73 | 0.05 |
| | cortical | venous diameter | -0.19 | 0.85 |
| histology | Parvalbumin GM | arterial density | 0.27 | 0.33 |
| | Nissl GM | arterial density | 0.42 | 0.08 |
| | Silver WM | arterial density | 0.49 | 0.05 |
| | Parvalbumin GM | venous density | 0.18 | 0.49 |
| | Nissl GM | venous density | 0.04 | 0.84 |
| | Silver WM | venous density | 0.43 | 0.08 |
| | Parvalbumin GM | arterial diameter | 0.19 | 0.48 |
| | Nissl GM | arterial diameter | 0.61 | **0.01** |
| | Silver WM | arterial diameter | 0.27 | 0.33 |
| | Parvalbumin GM | venous diameter | 0.04 | 0.84 |
| | Nissl GM | venous diameter | 0.25 | 0.36 |
| | Silver WM | venous diameter | 0.09 | 0.78 |

*Table E4: Spearman correlations between in-vivo vascular cerebellar measures and post-mortem cytoarchitecture and reference morphological measures.*
